# Improved genetic discovery and fine-mapping resolution through multivariate latent factor analysis of high-dimensional traits

**DOI:** 10.1101/2024.08.23.609452

**Authors:** Feng Zhou, William J Astle, Adam S Butterworth, Jennifer L Asimit

## Abstract

Genome-wide association studies (GWAS) of high-dimensional traits, such as molecular phenotypes or imaging features, often use univariate approaches, ignoring information from related traits. Biological mechanisms generating variation in high-dimensional traits can be captured parsimoniously through GWAS of a smaller number of latent factors from factor analysis. Here, we introduce a zero-correlation multi-trait fine-mapping approach, flashfmZero, for any number of latent factors. In our application to 25 latent factors derived from 99 blood cell traits in the INTERVAL cohort, we show how GWAS of latent factors enables detection of signals that have sub-threshold associations with several blood cell traits. FlashfmZero resulted in 99% credible sets with the same size or fewer variants than those for blood cell traits in 87% of our comparisons, and all latent trait fine-mapping credible sets were subsets of those from flashfmZero. These analysis techniques give enhanced power for discovery and fine-mapping for many traits.

## Introduction

Many genetic variants associated with disease risks or quantitative traits have been identified by genome-wide association studies (GWAS)[1]. There are many examples of pleiotropy amongst these findings, where a variant affects several traits, often by affecting a pathway upstream of multiple related traits[2]. When genetic variants affect a group of traits through a common pathway, methods that leverage the shared signal in the component of genetic variation common to all the traits, while accounting for residual correlation, are able to identify associated variants (multi-trait GWAS, e.g. MTAG[3]) and pinpoint causal variants (multi-trait fine-mapping, e.g. flashfm[4]) more powerfully than methods that analyse traits individually. Such approaches provide an efficient way to gain statistical power without increasing sample size. For example, multi-trait fine-mapping of four lipid traits in a GWAS of 6,407 Ugandan participants[5] gave similar results to single-trait fine-mapping in a meta-analysis of >125,000 African ancestry participants[6]. The *APOE* variant rs7412, an established causal variant for lipid levels, was not prioritised for high-density lipoprotein (HDL) cholesterol in the single-trait fine-mapping, but was prioritised by flashfm and confirmed as the probable causal variant by single-trait fine-mapping in the larger African ancestry meta-analysis.

Genetic studies of large numbers of high-dimensional traits, such as gene expression, protein or metabolite levels, often rely on univariate analyses, perhaps because most multi-trait GWAS methods are limited computationally to a handful of traits. Consequently, such studies do not leverage the information in association signals shared across the phenotypes. Some multi-trait GWAS methods use individual-level data to fit a multivariate linear model, jointly testing for association between a variant and each of multiple traits, but they are often limited computationally by the number of traits (e.g. GEMMA[7]). Summary-level (GWAS summary statistics) methods have the advantage that their computational efficiency does not depend on the sample size. They can be broadly partitioned into: (i) methods for joint modelling of the effect size estimates from multiple traits (e.g. MTAG[3]), which are not designed for high-dimensional phenotypes; (ii) approaches to reduce the dimension of the genome-wide joint distribution of the GWAS effect sizes from multiple traits through factor analysis (e.g. genomicSEM[8] or FactorGo[9]).

Rather than taking a dimension reduction approach to the distribution of effect sizes aggregated from multiple single-trait GWAS, we take a different perspective and investigate the GWAS of latent factors that underlie the original traits, through use of factor analysis (FA). FA captures the covariation between multiple traits by modelling them jointly as linear combinations of a set of common latent factors (plus independent error terms). Such latent factors can correspond to common sources of biological variation for which we can estimate GWAS summary statistics.

Current multi-trait fine-mapping methods that allow for multiple causal variants are not scalable to high-dimensional traits. Flashfm[4] multi-trait fine-mapping leverages information between traits in a Bayesian framework, giving improved resolution when traits share causal variant(s) and otherwise giving similar results to single-trait fine-mapping. Here, we extend flashfm to any number of latent factors by taking advantage of the zero correlation between the latent factors that we construct from FA with a varimax rotation. This modified version of flashfm rapidly fine-maps signals amongst multiple uncorrelated traits (or latent factors); to distinguish this new approach from the original flashfm, we call it flashfmZero.

To illustrate the performance of latent factor GWAS and single/multiple latent factor fine-mapping and to compare them respectively to univariate GWAS and univariate fine-mapping of multiple traits, we focus on 99 blood cell traits measured in the INTERVAL cohort of UK blood donors[10–12]. We cross-check the results of our analysis with fine-mapping performed in UK Biobank as part of a much larger study[13]. We show that fine-mapping of association signals using the latent factor phenotypes enables improved resolution over fine-mapping of the measured blood cell traits, with further gains through multi-trait fine-mapping of the latent factors by flashfmZero.

## Results

### Identification of latent factors underlying variation in blood cell traits

We used blood cell trait data from the extended complete blood count (CBC) reports generated using Sysmex XN haematology analysers in >45,000 generally healthy UK blood donors from the INTERVAL study[10–12] (Table S1). To identify groups of these blood cell traits that share common underlying latent factors, we applied factor analysis with the varimax rotation to data from 18,310 participants in INTERVAL with complete data (i.e. no missing measurements) for each of the 99 blood cell traits. We estimated 25 latent factors that are statistically uncorrelated (Methods).

We calculate scaled factor loadings *C_ij_*that indicate the contribution of each latent factor *j* to each blood cell trait *i*, i.e. the proportion of variance in blood cell trait *i* that is explained by latent factor *j*, relative to the total variance explained jointly by the 25 latent factors (Methods). In general, sets of blood cell traits that receive high ranking contributions to variance from the same latent factor belong to the same broad category of cell type (Figure 1, Table S2). For example, latent factor ML4 ranks highly amongst latent factors explaining variation in reticulocyte traits, but not in traits of other cell types. Likewise, ML8 ranks highly only in basophil traits, and ML5 ranks highly only in platelet traits.

**Figure 1.**
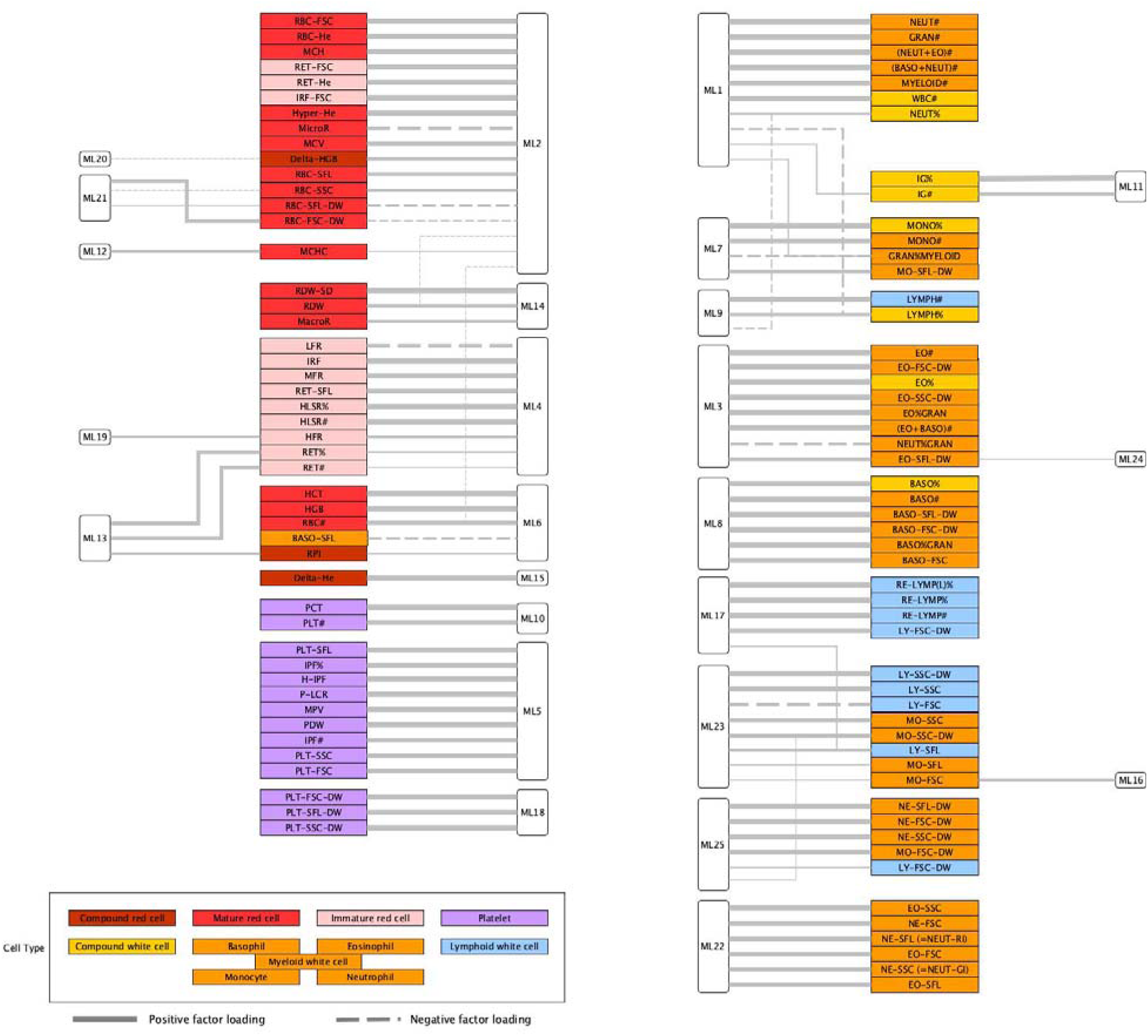
Latent factors partition blood cell traits grouped by broad cell-type into groups with common underlying variance generating mechanisms. A line between a latent factor (open rounded rectangles; e.g. ML11) and blood cell trait (coloured rectangles; e.g. IG% or IG#) indicates that the latent factor makes a contribution of at least 20% to the variance in the blood cell trait; line thickness is proportional to the corresponding C*_ij_*; solid lines indicate a positive factor loading and dashed lines indicate a negative factor loading. Blood cell traits are categorised by broad cell type, according to the colour-code in the legend. Compound red blood cell, mature red blood cell, and immature red blood cell traits are all red blood cell traits. Compound white cell, lymphocyte, eosinophil, monocyte, basophil, and neutrophil traits are all white blood cell traits. A compound red cell trait is a trait that depends on measurements of mature red blood cells and reticulocytes, while a compound white cell trait is a trait that depends on measurements of lymphoid and myeloid white cells. Standard abbreviations are given for each blood cell trait; the full names are available in Table S1. The source data are in Table S2 and further interpretations in Table S3.

We describe the principal effects of the latent factors in Table S3. In doing so, we consider the magnitude of the relative contributions that each makes to variation in the blood cell traits, as well as the directions of the factor loadings. For example, increased ML10 corresponds to increased platelet count without a change in platelet volume or other platelet characteristics. The volume of blood occupied by platelets (PCT) therefore goes up with ML10. Increased ML17 corresponds to more reactive lymphocytes, while increased ML23 corresponds to reduced average cell volume, increased average cellular complexity, increased variability in cellular complexity and increased average RNA content of both lymphocytes and monocytes.

Importantly, multiple latent factors make major contributions (i.e. C*_ij_*> 20%) to variation in some blood cell traits and there is not a simple one-to-one mapping between latent factors and blood cell traits. For example, latent factor ML2 — which varies closely with the mass of haemoglobin per red blood cell (MCH) — is a major contributor to variation in multiple red blood cell traits. It is responsible for 41% of the latent factor generated variation in RBC-SFL-DW and 96% of the latent factor generated variation in RBC-FSC. On the other hand, although ML21 — a latent factor that principally affects the distribution width of the mass of haemoglobin in red cells (RBC-FSC-DW) — is responsible for 25% of the latent factor generated variation in RBC-SFL-DW, it contributes very little to variation in RBC-FSC. Notably, blood cell traits from the same broad category do not necessarily have the same primary contributing factor. For instance, ML1 — which varies closely with neutrophil count (NEUT#) — is the primary factor contributing to seven white cell traits that are direct functions of NEUT#, for six of which it contributes more than 83% of the total latent factor generated variance. On the other hand, ML22 makes the principal contribution to variation in average neutrophil volume (NEUT-FSC, 91%), average neutrophil granularity/complexity (NEUT-SSC, 69%) and average neutrophil nucleic acid content (NEUT-SFL, 88%). ML22 also makes the principal contribution to average eosinophil volume (EO-FSC, 70%), average eosinophil granularity/complexity (EO-SSC, 93%) and average eosinophil nucleic acid content (EO-SFL, 63%), suggesting the latent factor captures a biological mechanism common to the two types of granulocytes.

### GWAS of latent factors identifies additional association signals over blood cell trait GWAS

For each of the 25 latent factors and 99 blood cell traits, we conducted a GWAS using data from the 18,310 participants who contributed to the factor analysis. To identify genetic association signals discovered by the latent factor GWAS but not the blood cell trait GWAS, we first selected the variants associated (*P* < 5×10^-8^) with each latent factor and clumped them by linkage-disequilibrium (LD; *r*^2^ > 0.6). We then identified the blood cell traits that receive a contribution of at least 1% from the latent factor and examined the blood cell trait *p*-values for association with the lead variants (those with the smallest *p*-value) from each of the latent factor clumps. Out of 3,399 lead variant associations, 3,036 had a genome-wide significant (*P* < 5×10^-8^) association with a connected blood cell trait, 211 had evidence for association at a suggestive significance threshold (i.e. 5×10^-8^ < *P* < 1×10^-6^) and 152 did not have evidence for association at a suggestive significance threshold (*P* < 1×10^-6^) at any of the connected blood cell traits (Table S4).

Next, to explore the signals across latent factors and blood cell traits, we formed LD clumps (*r^2^* > 0.6) of the set of variants with a genome-wide significant association with at least one of the 99 blood cell traits; separately, we formed clumps of the set of variants with a genome-wide significant association with at least one of the 25 latent factors. We assumed each clump to represent a distinct association signal and considered a signal identified by the blood cell traits to have been identified by the latent factors if any variant in the corresponding blood cell trait clump exhibited a genome-wide significant association with a latent trait. Symmetrically, we considered a latent factor signal to have been identified by the blood cell traits if any variant in the corresponding clump exhibited a genome-wide significant association with a blood cell trait. As expected, we found that blood cell trait clumps that are significantly associated with multiple blood cell traits are more likely to be significantly associated with a latent factor (Armitage test for trend *P*<10^-25^); 67% (98/146) of the clumps associated with exactly two blood cell traits and 86% (102/119) of the variants associated with exactly three blood cell traits were also associated with a latent factor (Figure 2, Table S5). In contrast, 32% (96/301) of clumps associated with just one blood cell trait were also associated with a latent factor.

**Figure 2.**
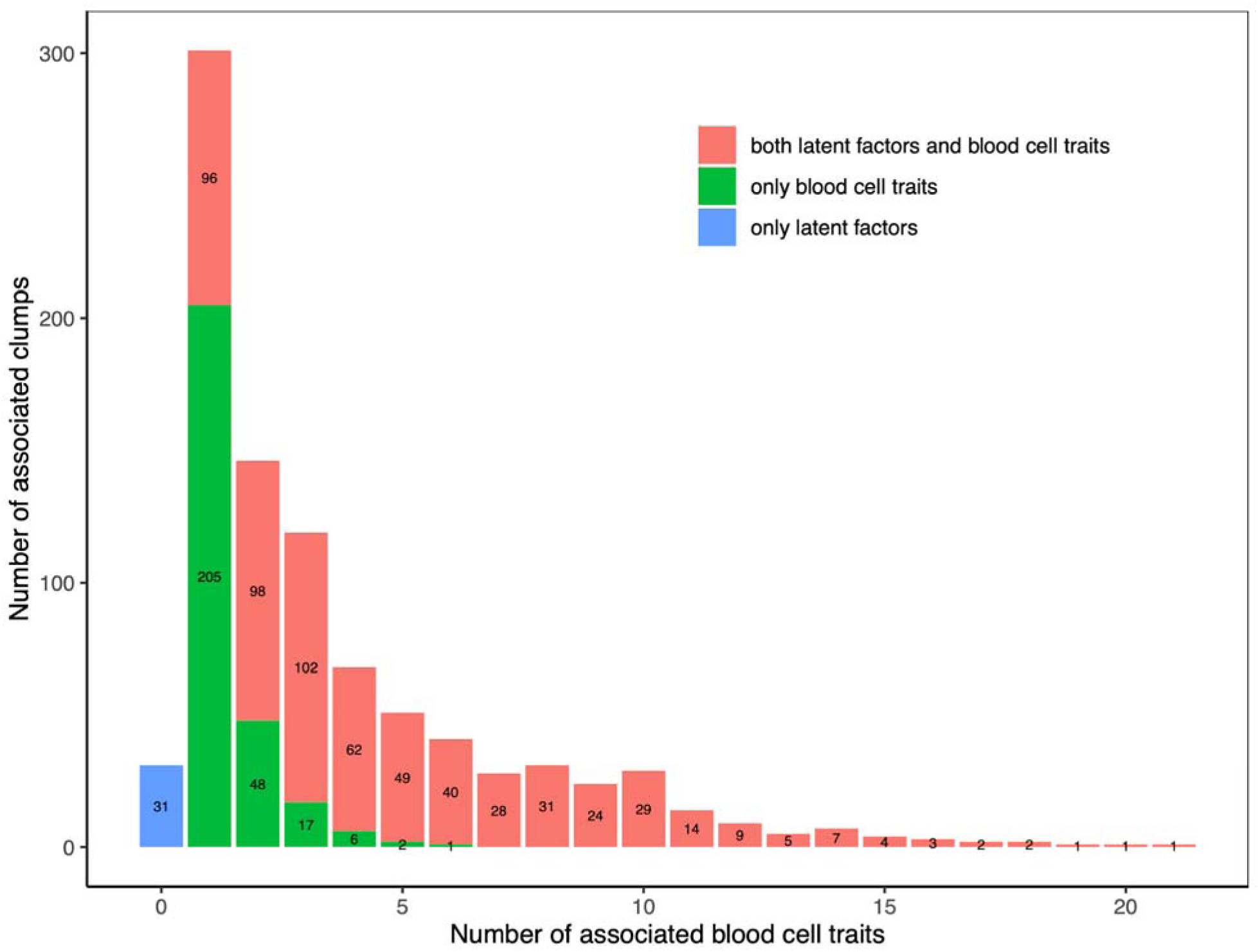
Clumps that are associated with multiple blood cell traits are more likely to be associated with a latent factor. This stacked barplot counts both blood-cell-trait associated variant clumps and latent factor associated variant clumps (MAF > 0.003, LD clumped *r^2^* > 0.6). The *x*-axis indicates the number of distinct blood cell traits associated (*P* < 5×10^-8^) with at least one variant in a clump. The *y*-axis indicates frequency. Red bars count blood cell trait signals shared with at least one latent factor, while green bars count blood cell trait clumps not associated with latent factors. The blue bar shows that 31 clumps are associated with a latent factor but not a blood cell trait. The source data are given in Table S5(B).

The advantage of factor analysis is illustrated by the 31 clumps that do not exhibit genome-wide significant evidence for association with a blood cell trait, but that are significantly associated with a latent factor because they are moderately associated with several blood cell traits. For example, we found an association between ML8 and rs9310935 near *IL5RA* (per allele effect size estimate = -0.058SD, 95% confidence interval (-0.078SD, -0.037SD), *P* = 3.3×10^-8^) (Figure 3), a variant previously detected in a multi-ancestry meta-analysis for basophil count[14], but which did not reach genome-wide significance in our single blood cell trait GWAS nor in substantially larger GWAS of European ancestry participants[10,13]. However, we did find moderate-to-weak evidence for association between rs9310935 and four basophil-related traits to which ML8 contributes variance (Figure 3, Table S4). Several other ML8-associated variants in the *IL5RA* region were associated with white blood cell traits in our analysis, as well as in previous studies (Figure 3, Figure S1) [11]. rs9310935 remains significantly associated with ML8 when conditioning on the published lead variants, suggesting that its association signal is distinct from those previously identified (Figure 3, Figure S1).

**Figure 3.**
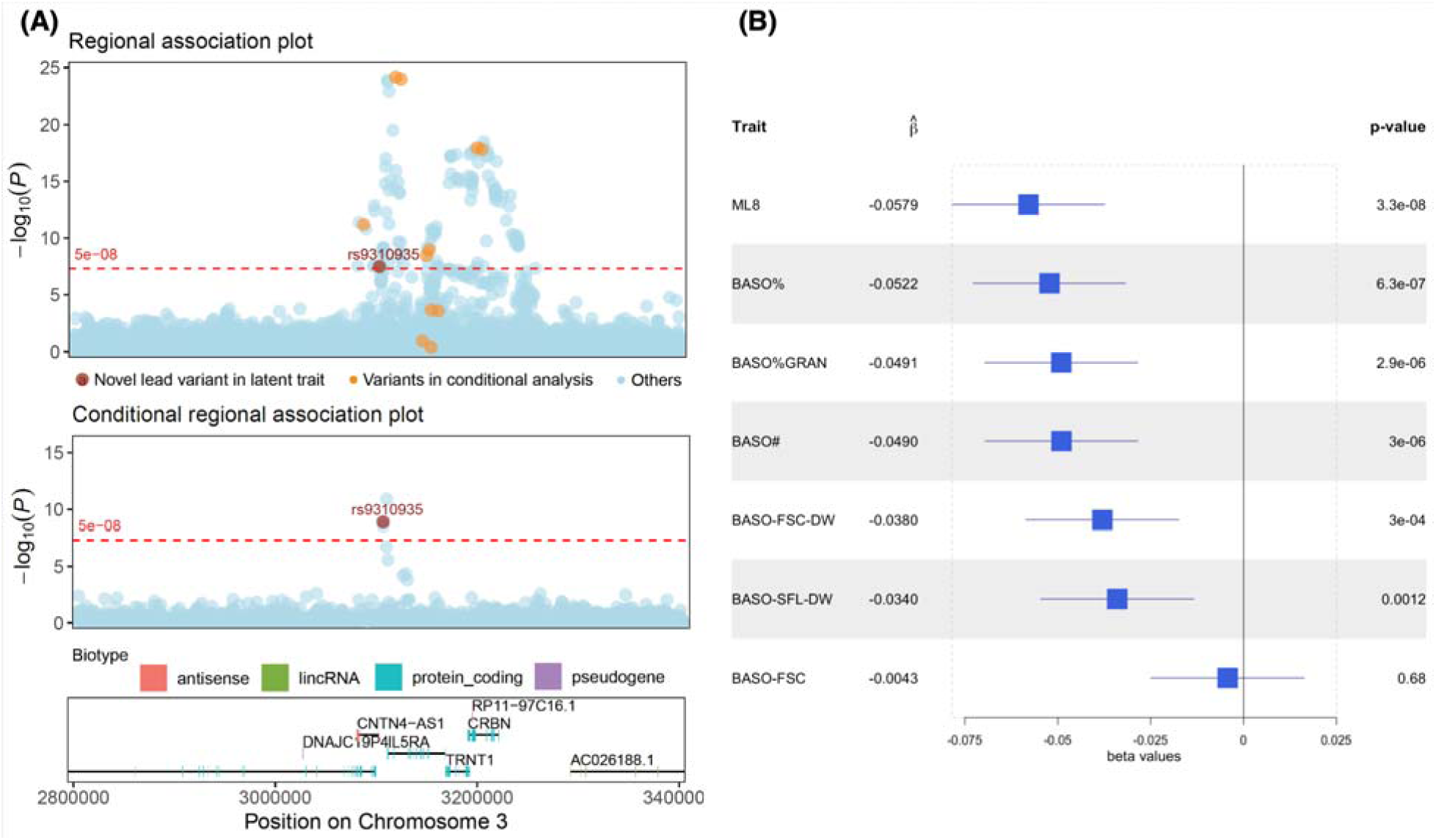
Basophil-related latent factor ML8 is associated with rs9310935, which exhibits moderate evidence for association with multiple basophil-related traits. (A) Regional association plot for ML8 (top panel), highlighting rs9310935 and conditional regional association plot for ML8 (middle panel), conditioned on, from top to bottom, rs3217673, rs163546, rs1669340, rs17027750, rs163563, rs3856850, rs334782, rs1695315, rs10212483, rs13097407, rs6787336, which are lead SNPs for basophil-related traits from previous publications (rs163546 (BASO%GRAN), rs3856850/rs10212483/rs163563/rs17027750 (BASO%), rs13097407 (EO%), rs1695315 (EO#), rs334782 (BASO# and BASO-FSC-DW), rs1669340 (BASO-SFL-DW and BASO-FSC-DW), rs3217673 (BASO%, BASO-SFL-DW and BASO-FSC-DW) and rs6787336 ((EO+BASO)#)). See Figure S1 for a labelling of the conditioned variants highlighted in (A); (B) Forest plot showing estimates of the effect size (change in mean per copy of alternate allele) and corresponding 95% confidence intervals of the associations between rs9310935 and ML8 and the six basophil traits to which ML8 contributes substantial variance.

**Figure 4.**
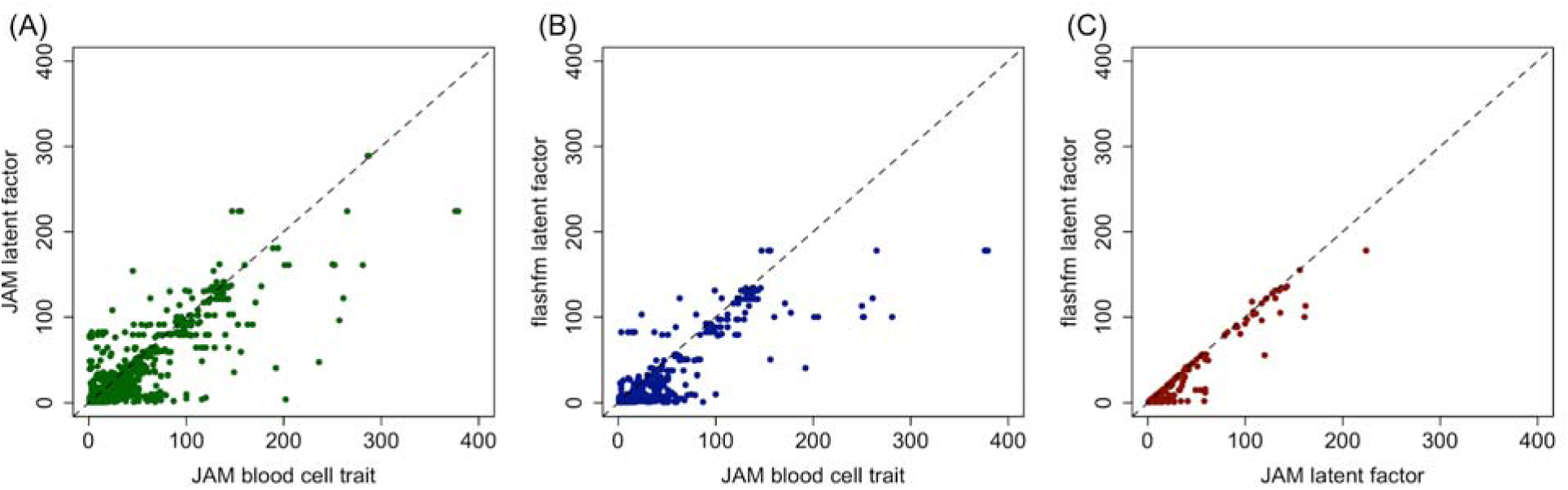
Latent factor fine-mapping yields smaller 99% credible sets than raw blood cell trait fine-mapping, with the largest gain in resolution from joint fine-mapping of multiple latent factors. ‘JAM latent factor’ tends to improve resolution over ‘JAM blood cell trait’, and further resolution gains are observed from ‘flashfm latent factor’ over ‘JAM latent factor’. The source data are given in Table S7.

Despite analysing a relatively small sample (18,310 participants) compared to previously published GWAS of blood cell traits, we identified five distinct genome-wide significant associations with the latent factors after conditioning on the 3,559 lead variants from recent large GWAS of complete blood cell traits[11] and Sysmex extended blood count traits[10]. None of the five associated variants showed genome-wide significant evidence for association in our univariate analyses of the blood cell traits in the same participants (Methods). Four of the variants exhibiting novel associations were common (MAF>0.01) and two were low-frequency (0.003 < MAF < 0.01) (Table 1, Table S6).

**Table 1.** Latent factors allow the identification of associations that are not detectable in GWAS of individual blood cell traits. . Conditional *p*-values are calculated from association analyses conditioning on the lead variants that have been previously identified by large GWAS of blood cell traits[10,11]. Further details are in Table S6. Variant physical positions are given in GRCh37 coordinates.

| Variant (chr:bp, rsID) | MAF | Nearest gene | Latent Factor (cell type) | Conditional $p$-value | Conditional $\beta$ (standard error) | Blood cell traits min $p$-value (trait name) |
| --- | --- | --- | --- | --- | --- | --- |
| 3:3101825 (rs9310935) | 0.4762 | *CNTN4* | ML8 (basophils) | $6.5 \times 10^{-9}$ | -0.0630 (0.0104) | $2.3 \times 10^{-7}$ (BASO%) |
| 5:40975803 (rs540446526) | 0.0036 | *C7* | ML18 (platelet) | $1.0 \times 10^{-8}$ | -0.5132 (0.0901) | $7.0 \times 10^{-7}$ (PLT-FSC-DW) |
| 5:41018263 (rs559314725) | 0.0031 | *MROH2B* | ML18 (platelet) | $2.2 \times 10^{-8}$ | -0.5402 (0.0971) | $4.8 \times 10^{-7}$ (PLT-FSC-DW) |
| 18:23024328 (rs72878322) | 0.3587 | *ZNF521* | ML8 (basophils) | $1.8 \times 10^{-8}$ | -0.0624 (0.0112) | $2.8 \times 10^{-7}$ (BASO%) |
| 20:54884793 (rs6064377) | 0.3277 | *FAM210B / MC3R* | ML6 (mature red blood cells) | $4.4 \times 10^{-8}$ | -0.0618 (0.0112) | $1.8 \times 10^{-7}$ (HCT) |
| 21:23426550 (rs117617749) | 0.0642 | AP000472.2 | ML6 (mature red blood cells) | $4.1 \times 10^{-8}$ | 0.1172 (0.0214) | $2.9 \times 10^{-7}$ (RBC#) |

One of the common variants (rs6064377, MAF=0.33) was associated with the latent factor ML6 (per allele effect size estimate = -0.0609SD, 95% confidence interval (-0.083SD, -0.039SD), *P* = 4.6×10^-8^), variation in which causes a change in haemoglobin (HGB) concentration and hematocrit (HCT) mediated by a change in red blood cell count (RBC#) while mean red corpuscular volume (MCV) and mean corpuscular haemoglobin (MCH) remain constant. The variant exhibited moderate evidence for association with HGB, HCT and RBC, with *p*-values between 1.5×10^-7^ and 1.3×10^-5^ (Figure S2). rs6064377 lies near the gene *FAM210B*, which has been shown to play a role in erythropoiesis, consistent with the associations discovered with red blood cell traits[15].

### Fine-mapping resolution gains are highest for joint latent factor fine-mapping

We considered 217 genomic regions that contain genetic association signals (*P* < 5×10^-8^) with any raw blood cell trait at least 20% of the variance of which is explained by a latent factor with a signal in the same region (Methods). Within each region, we applied JAMdynamic single-trait fine-mapping to each latent factor (‘JAM latent factor’) with a suggestive association signal (*P* < 1×10^-6^) in the region and to all raw blood cell traits (‘JAM blood cell trait’) that receive a contribution of at least 20% from these latent factors and have a suggestive association signal in the region. We also applied multi-trait fine-mapping to the latent factors (‘flashfm latent factor’) with flashfmZero (Methods).

‘JAM latent factor’ had a 99% credible set (CS99) that was smaller or equal to that of ‘JAM blood cell trait’ in 76% of the comparisons (and strictly smaller in 58% of the comparisons). ‘Flashfm latent factor’ had a CS99 that was smaller or equal to that of ‘JAM blood cell trait’ in 87% of the comparisons (and strictly smaller in 71% of the comparisons). When comparing the single and multi-trait latent factor approaches, we found the CS99 from fine-mapping by ‘flashfm latent factor’ to be smaller or equal to that from ‘JAM latent factor’ in 97% of the comparisons (and strictly smaller in 45% of comparisons).

To assess the accuracy of our latent factor fine-mapping results, we turned to the marginal posterior probability (MPP) of causality for a variant. We compared the variants identified as causal with high-confidence (MPP>0.90) by our analyses in the INTERVAL sample of approximately 18k individuals with those identified (MPP>0.95) in the UK Biobank sample of approximately 500k individuals[13]; we allowed a lower prioritisation threshold in our comparatively smaller analysis, although note that the majority of our high-confidence variants do satisfy MPP>0.95 (Table S7).

For this comparison we focus on 36 regions that met two conditions: (i) contained a variant identified with high confidence as causal by fine-mapping with either ‘JAM blood cell trait’, ‘JAM latent factor’, or ‘flashfm latent factor’ in the INTERVAL analysis (MPP>0.90), as well as by fine-mapping with FINEMAP[16] in the UK Biobank analysis (PP>0.95)[13]; (ii) the causal association identified in the INTERVAL fine-mapping was with one of the 29 ‘classical’ CBC traits analysed in the UK Biobank study or with a latent factor connected to such a trait (Methods). In 69% (25/36) of the regions, at least one of the high-confidence variants identified by blood cell trait and latent factor approaches matched those identified in UK Biobank (Table S7). In an additional 11% (4/36) of the regions, the high confidence variants identified by the latent factor approaches matched those identified in UK Biobank but the fine-mapping of the original blood cell traits did not identify any high-confidence causal variants.

Amongst the 25 regions in which the fine-mapped variants from both the blood trait and latent factor approaches match those identified in UK Biobank we found 9 regions for which latent factor fine-mapping enabled identification of the likely causal variant with high confidence for more traits than blood cell trait fine-mapping. For example, in a region containing *PIEZO1* (a gene with a primary role in blood vessel formation and vascular structure[17]), there were four correlated variants (*r*^2^ > 0.8) in the CS99 for the mature red cell trait mean corpuscular haemoglobin concentration (MCHC), of which rs861400 and rs551118 had the highest (0.69) and second highest (0.27) MPP, respectively (Figure S4). However, rs551118 had the highest MPP for the immature red cell traits reticulocyte count (RET#; MPP = 0.94; CS99 size = 4), reticulocyte percentage (RET%; MPP = 0.88; CS99 size = 8) and reticulocyte production index (RPI; MPP=0.56; CS99 sizes = 8). The primary contributor to MCHC is the latent factor ML12, while the primary contributor to RET#, RET%, and RPI is the latent factor ML13. Our joint fine-mapping of the latent factors led to high-confidence that rs551118 is causally associated with both ML12 (MPP=0.97) and ML13 (MPP=0.96). Our results suggest that rs551118 is the likely causal variant for MCHC, RET, RET%, and RPI, which are the traits with high contributions from ML12 and ML13.

There were four regions in which only the joint fine-mapping of latent factors was able to prioritise high confidence variants to match those identified in the UK Biobank. In one of these regions, our joint fine-mapping prioritised rs1175550 an intronic variant of *SMIM1* (a regulator of red blood cell formation and the gene encoding the antigen underlying the Vel blood group[18]), for three latent factors (ML4, ML12, ML14) that are all related to red blood cell traits (Figure 5). This result was validated by the UK Biobank fine-mapping which identified rs1175550 as a high-confidence variant for nine red blood cell traits (e.g. HGB, RBC#, MCHC, RET#). It is also supported by published data showing rs1175550 to be an eQTL for *SMIM1* and a modulator of Vel blood group antigen expression[19]. The CS99 from the joint latent factor fine-mapping contained a single variant, a noticeable improvement over the results of the fine-mapping of the original blood cell traits, for which the CS99s — all containing rs1175550 — ranged in size from 30 to 58 (Figure 6). rs1175550 had the highest MPP (0.24–0.59) in the univariate fine-mappings of the associations with HLSR#, HLSR%,MFR and MCHC, and the second highest MPP in the fine-mapping of the associations with IFR and LFR (MPP≈0.20), for which the highest confidence variant (rs1175549, MPP≈0.26) is in high LD (*r*^2^=0.83) with rs1175550. rs1175550 was also ranked second (MPP=0.12) — after rs7513053 with which it exhibits moderate LD (*r*^2^=0.69, MPP=0.44) — in the fine-mapping for RDW-SD. It is only ranked eighth (in a CS99 of 46 variants) in the fine-mapping for RET-SFL and is only in moderate LD (*r*^2^=0.47) with the top variant (rs1175548, MPP=0.33). Single-trait fine-mapping of latent factors improves resolution over that of the blood cell traits, with CS99 sizes ranging from 5 to 27; rs1175550 had highest MPP for ML4 (MPP=0.42) and ML12 (MPP=0.92) and is ranked second for ML14 (MPP=0.18), though it is in LD with the top variant, rs7513053 (*r*^2^=0.69, MPP=0.33). Multi-trait fine-mapping further refines the CS99 for the three latent traits, with rs1175550 having MPP > 0.99 (Figure 6).

**Figure 5.**
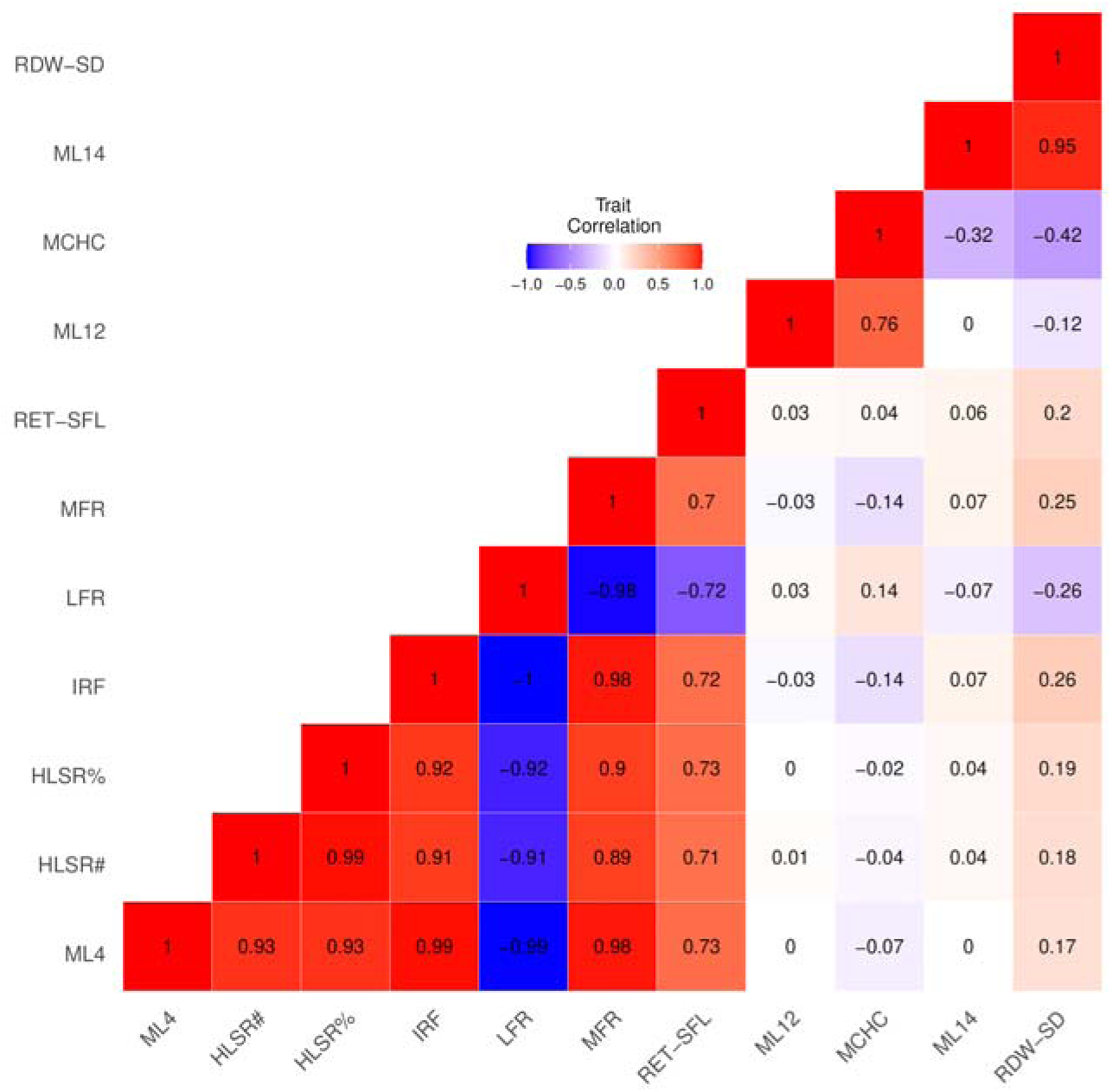
The correlation between the latent factors and blood cell traits exhibiting associations in the *SMIM1* region. There are high correlations between each latent factor and the blood cell traits to which they contribute substantial variance, and high correlations amongst blood cell traits with contributions from a common latent factor. The three latent factors only contribute substantial variance to red blood cell traits — ML4 contributes to six traits, ML12 contributes to MCHC, and ML14 contributes to RDW, as indicated by the correlation blocks. Similar plots for the phenotypes exhibiting genetic associations in the regions containing *PIEZO1* and the *TMCC2* are shown in Figure S3 and Figure S5.

**Figure 6.**
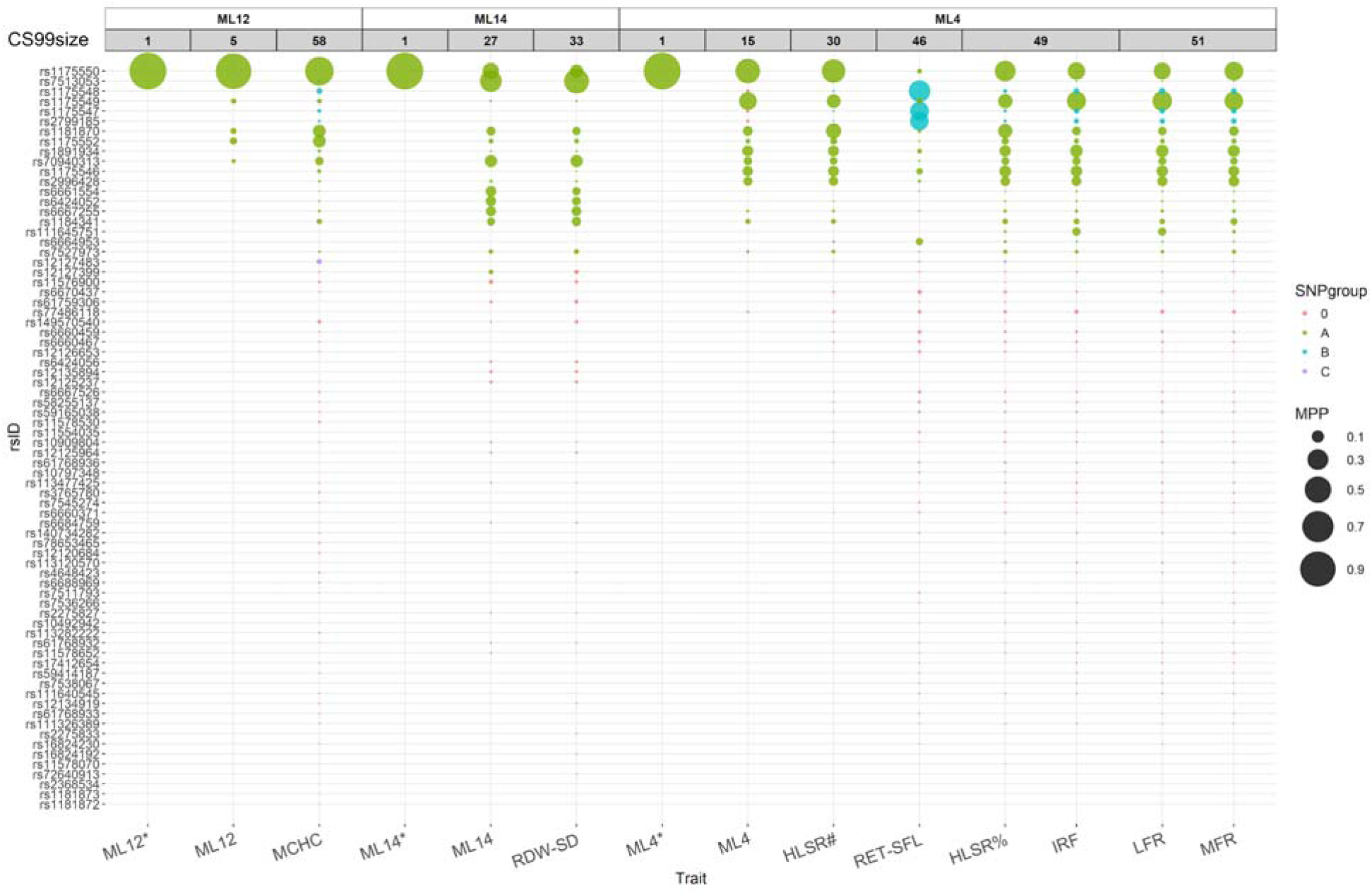
In the region containing *SMIM1*, the 99% credible sets (CS99) derived from latent factor multi-trait fine-mapping contain a single variant, refining those derived from univariate fine-mapping by latent factors, which in turn refine those derived from univariate fine-mapping of blood cell traits. The variants indicated by rsID on the *y*-axis belong to at least one CS99 from the various fine-mappings. Each column corresponds to the CS99 from the fine-mapping indicated on the *x*-axis. The CS99 for the univariate latent factor fine-mappings are denoted by the latent factor name (e.g. ML12). The CS99 for multi-trait fine-mapping of latent factors are denoted by the latent factor name appended with an asterisk (e.g. ML12*). The coloured circles indicate which variants are members of the CS99 for the listed trait.The area of each circle is proportional to the marginal posterior probability (MPP) that the variant is causally associated with the trait. The colours of the circles indicate groups of variants (with MPP > 0.01) in high LD (*r^2^* > 0.8)as calculated by the fine-mapping method. Columns are grouped (open boxes top row) according to whether or not the latent factors contribute to the blood cell traits. Within each group, the columns are ordered by the sizes of the corresponding CS99s (e.g. ML12* has a CS99 of size 1, ML12 has a CS99 of size 5, and MCHC has 58 variants in its CS99).Similar plots for the *PIEZO1* and *TMCC2* regions are shown in Figure S4 and Figure S6.

Where latent factors are not biologically related, the multi-trait fine-mapping results are identical to the fine-mapping results for the individual latent factors. For example, in a region containing *TMCC2,* we fine-mapped signals from 12 blood cell traits, of which nine are platelet traits and three are basophil traits (Figure S5). No variants were prioritised with high confidence by the fine-mapping of the individual blood traits. The CS99 sizes ranged from 16 to 52 for the platelet traits and 18 to 19 for the basophil traits (Figure S6). The nine platelet traits are linked to ML5 for which fine-mapping results in a CS99 containing eight variants. Likewise, the three basophil traits are linked to ML8, for which the size of CS99 was reduced to 12. The CS99 of the basophil and platelet traits do not overlap, suggesting that they are unlikely to share any causal variants in this region, and hence our flashfm (multi-trait) latent factor results were identical to those from individual (single-trait) latent factor fine-mapping (Figure S6).

## Discussion

Using blood cell traits as a model, we have shown that where multiple phenotypes capturing common biological variation have been measured, genetic association analysis of latent variables derived by factor analysis can provide a complementary approach to univariate genetic association analyses. The factor analysis approach has three main advantages. Firstly, the latent factors help to identify groups of measured traits in which variation is generated by common underlying biological mechanisms. Consequently, analysis of these latent factors leads to inferences about groups of traits that share the same underlying factors. Secondly, GWAS of latent factors can boost power when a variant exhibits only moderate evidence of association with each of multiple measured traits, enabling detection of signals that are missed by GWAS of the individual measured l traits. Finally, in the context of studies measuring hundreds or thousands of phenotypes the use of factor analysis to construct latent factors could be favourable from the perspective of green computing[20], either as a substitute for univariate analysis of the measured traits or as a pre-filtering step to identify variants unlikely to be associated in the univariate GWAS.

The example of *SMIM1* demonstrates that multi-trait fine-mapping of latent factors using flashfmZero, a zero-correlation version of flashfm which can be applied to any number of traits, can significantly improve resolution. This is because orthogonal latent factors may share causal variants if they capture aspects of a common biological process, despite the fact that they are, by mathematical definition, uncorrelated. However, we note that when the latent factors are not biologically related, it is less likely that they will share causal variants. In such instances, multi-trait latent factor fine-mapping will give similar results to univariate latent factor fine-mapping, though there will often be resolution gains over univariate fine-mapping of the measured traits.

Latent factor approaches are limited to analyses where individual-level data are available on the traits and genotypes. The trait measurements are required to calculate the latent factors and the genotypes are needed to perform the genetic association analysis and to estimate LD for fine-mapping. It is possible to derive factor loadings from a trait correlation matrix, but individual-level trait data are required to compute factor scores. Factor loadings are useful to determine which latent factors contribute to the measured traits, but the genetic association analysis requires access to the factor scores.

The computation of the latent factors scores for the participants in a study cohort requires complete data, i.e. any participant with a missing measurement for any trait must be excluded. Consequently, the sample available for our analysis was a little over 18,000 of the approximately 40,000 individuals in the INTERVAL cohort. Given the extent of the missing data and because measurements from multiple traits were often missing in the same individuals due to a period in which a haematology analyser was faulty (Methods), we did not explore imputation of the missing trait measurements. In any case, our aim was to demonstrate the advantages of factor analysis approaches by comparison with univariate approaches in a sample of a given size. Nevertheless, there may be contexts in which the imputation of missing data could allow association analyses with larger sample sizes and improve power, as demonstrated by increased genetic discovery in UK Biobank GWAS, where deep-learning based trait imputation was implemented using AutoComplete[21].

Our analyses have illustrated the value of latent factor GWAS, with clear gains in fine-mapping, especially when signals from multiple latent traits are jointly fine-mapped. Further gains in fine-mapping resolution could be attained by incorporating functional annotations in the prior probabilities, an approach taken in PAINTOR[22] and PolyFun[23].

## Methods

### Experimental model and subject details

#### INTERVAL cohort

INTERVAL is a cohort of 48,725 generally healthy adult blood donors recruited through NHS Blood and Transplant, the English Blood Service, between 2012 and 2014[12,24]. The cohort was originally established for a clinical trial to assess the effect of variation in inter-donation time intervals on the health of blood donors[25]. The study was approved by the Cambridge East Research Ethics Committee and informed consent was obtained from all participants during recruitment.

Participants were genotyped by Affymetrix (Santa Clara, Ca, USA) with the UK Biobank Axiom array using DNA extracted from buffy coat by LGC Genomics (UK). Standard Affymetrix quality control (QC) procedures were applied to the resulting data, which excluded genotyping probes with low signal intensity, samples with low call rates and variants with low call rates or low confidence calls. Further QC procedures were applied — to the full dataset and within each genotyping batch — to remove rare variants, multiallelic variants, variants with a poor call rate, variants out of HWE and variants exhibiting allele frequency variability across batches. Variants were pruned to ensure no pair exhibited strong LD. Samples exhibiting evidence for contamination, excess heterozygosity, non-European ancestry or discordance of phenotypic and genotypic sex were removed. Subsequently, haplotype phases were imputed using SHAPEIT3 and missing genotypes were imputed from the 1000 Genomes Phase 3-UK10K reference panel using the PBWT[26,27].

Extended complete blood counts (CBCs) were measured from EDTA treated blood samples taken from INTERVAL participants using two Sysmex XN haematology analysers at UK Biocentre (Stockport, UK). Because a flow-cytometry channel of one instrument was misconfigured during the first 90 days of the study, data for some platelet variables are missing for some participants. The extended CBC produced by the Sysmex instrument measures various properties of the peripheral blood, including haemoglobin levels and properties of reticulocytes, mature red cells, platelets, neutrophils, eosinophils, basophils, monocytes and lymphocytes. These properties include cell concentrations, measures of cell maturity, properties of cell volume distributions and properties of the distributions of cell fluorescence and cell side-scatter measured by flow-cytometry.

Each variable in the CBC was adjusted to remove variance explained by technical covariables including, the identity of the measuring instrument, the age of the blood sample at the time of measurement, the time of day of the measurement, time dependent instrumental drift and instrument recalibration events. Measurements taken more than 36 hours after venipuncture were excluded. Subsequently, we adjusted each variable to remove variance explained by sex, menopausal status, age, smoking habits, drinking habits, log-height and log-weight. Finally, measurements that were outliers in univariate and cell-type specific multivariate distributions were removed. The phenotypes were then rank inverse normalised.

More detailed descriptions of the QC procedures applied to the genotype and phenotype data are given in [10] and [11].

### Method details

#### Factor analysis of quantitative traits

A trait correlation matrix is sufficient to construct latent factors by computing their loadings and to quantify the contribution of each factor (re-scaled factor loadings) to each original raw trait. However, our objective is not only to compute the loadings, but also to compute the values of the latent factors for each individual (i.e. the factor scores). This requires individual-level data. Let *L_ij_* be the factor loading of latent factor *j* (*j=1,…,L*) for raw trait *i* (*i=1,…T*). We define the contribution of latent factor *j* to raw trait *i* by 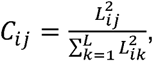, to aid in mapping the contributions of the latent factors back to each raw trait.

We only used participants that have measurements for all the measured traits, so that factor scores could be computed for each individual from the factor loadings and the measured traits. The application of imputation approaches such as Multivariate Imputation by Chained Equations (MICE)[28] was inappropriate, because the measurements were not missing independently by trait; subsets of individuals were missing certain platelet measurements, as described above in the INTERVAL cohort section. Consequently, rather than introducing noise through poor quality imputation, we opted to reduce the sample size.

The final sample size was 18,310. Each participant had a measurement for each of the 99 raw blood cell traits. We applied factor analysis in R using the “fa()” function in the psych library[29], with the arguments fm=”ml”, for a maximum likelihood factor analysis and rotate=”varimax”. The varimax rotation preserves the orthogonality of latent factors (factor scores), so that they are uncorrelated. A scree plot (using “fa.parallel()” in the psych package) indicated that 25 latent factors was an optimal choice.

#### GWAS and conditional analyses

Each of the 25 latent factors and 99 blood cell traits was tested for genetic associations within the sample of 18,310 individuals from the INTERVAL cohort using BOLT-LMM[30] with the following covariates: dummy variables indicating the donor clinic at which the blood sample was taken and the score vectors corresponding to the leading ten principal components of genetic variation in the study sample. This follows the approach taken in a previous large-scale GWAS of blood cell traits (that included the INTERVAL cohort) in 173,480 European descent individuals[11] and a GWAS of flow-cytometry derived (Sysmex) blood cell traits in 41,515 INTERVAL cohort participants[10]. All the traits were inverse normal-rank transformed prior to running BOLT-LMM.

To identify potentially novel association signals in our latent factor GWAS of 18,310 individuals, we conditioned on all the lead variants identified in the previously published large-scale GWAS of common blood cell traits[11] and the GWAS of Sysmex blood cell traits[10].

We obtained a list of unique variants that are genome-wide significant for any of the 99 blood cell traits, through LD clumping (*r^2^* > 0.6) on the merged list of associated variants. Then, at each unique variant we recorded the number of blood cell traits that were associated with the variant; if the variant was not associated with a blood cell trait, but it had a tag variant (in the same clump) that was associated, the trait was enumerated. To identify unique variants missed by blood cell traits, we enumerated the unique independent genome-wide significant variants obtained only by latent factors, based on LD clumping (*r^2^* > 0.6) of their associated variants, allowing for the variant or one of its tag variants to be associated.

#### Fine-mapping

To investigate gains from fine-mapping association signals using latent factors that are uncorrelated by construction, over fine-mapping association signals using a larger number of correlated traits, we constructed regions based on the latent factor association signals. For each latent factor, we used distance-based clumping to identify lead SNPs with a distance of at least 250kb, which were then centred ±250kb to form regions. Regions that overlapped amongst traits were merged. We further expanded our regions by integrating them with those from fine-mapping signals from 29 blood cell traits in UK Biobank[13] and merging any that overlapped, so that our regions contained those used in the UK Biobank fine-mapping. This resulted in 217 regions with lengths ranging from 500,000bp to 2,996,725bp.

Within these regions, we fine-mapped suggestive association signals (P<1×10^-6^, MAF>0.005) with all the latent factors and with all the blood cell traits that have a contribution of at least 20% from these latent factors. Single-trait fine-mapping of latent factors and blood cell traits was carried out with JAMdynamic[31], which is an extension of JAM[32] that dynamically selects the maximum number of causal variants based on the data. When multiple latent factors had a signal in a region, we also used our zero-correlation version of flashfm, as available in the wrapper function FLASHFMZEROwithJAMd(https://jennasimit.github.io/flashfmZero/).

For both methods, we used an LD matrix calculated from the 18,310 participants in the INTERVAL cohort that contributed to the GWAS. In particular, we used best-guess genotypes with a certainty threshold of 0.2, such that the genotype at a variant took on values 0,1, or 2 if their dosage was within 0.2 of the respective value; otherwise, the genotype was coded as NA in the correlation calculation. For each variant, we calculated the proportion of individuals with non-missing best-guess genotypes, and excluded any variants that had a non-missing proportion below 80%.

In our comparisons of the fine mapping resolution of the three approaches: (i) ‘JAM blood cell trait’ (JAMdynamic on each blood cell trait); (ii) ‘JAM latent factor’ (JAMdynamic on each latent factor); (iii) ‘flashfm latent factor’ (flashfmZero on each set of latent factors), we considered the CS99 size and variants with PP>0.90, matching on traits. That is, when comparing blood cell trait results to latent factor results, we match each blood cell trait to the latent factor that is the highest contributor to it.

We considered a variant to be a high-confidence causal variant if it had PP>0.90 and cross-checked our results with the high-confidence causal variants (PP>0.95) from the UK Biobank analysis[13] for validation; as our sample size is substantially smaller than that of UK Biobank, we used a slightly lower threshold when defining high-confidence. We identified 53 regions where a high-confidence variant was detected by either single-trait or multi-trait fine-mapping of the latent factor association signals and also detected in UK Biobank. Amongst these 53 regions, 17 regions are not comparable because our analyses included only latent factors that are linked to extended Sysmex traits, whereas the UK Biobank analyses did not include all the extended Sysmex traits. Therefore, we focused on 36 regions in cross-checking our latent factor fine-mapping results with those of the UK Biobank study. We also note that there were 9 regions where no high-confidence variants were identified by our latent factor analyses, but there was prioritisation by ‘JAM blood cell trait’ — in 5 of these regions there was alignment with the UK Biobank results and in the remaining 4 regions there was not an exact match in the high-confidence variants selected in INTERVAL and UK Biobank (Table S7).

#### Multi-trait fine-mapping with flashfmZero

The multi-trait fine-mapping method, flashfm[4], leverages information between traits while allowing for multiple causal variants that are not necessarily shared between traits. It is flexible to missing trait measurements. When there are shared causal variants, flashfm has been shown to improve fine-mapping resolution and increase the number of high-confidence variants, compared to single-trait fine-mapping[4][6]. Otherwise, it gives comparable results to single-trait fine-mapping.

Flashfm requires the trait correlation matrix and is currently limited to six traits at most. However, we take advantage of the uncorrelated latent factors that result from using a varimax rotation, resulting in a diagonal correlation matrix. Under this condition, we have extended flashfm to multiple-trait fine-mapping of an unlimited number of (uncorrelated) latent factors. We call this extended method flashfmZero.

For *M* uncorrelated traits, the joint Bayes’ factor *BF^M^* can be expressed as the product of the marginal trait *BF_j_, j=1,…M.* Without loss of generality, the next steps focus on *M*=2 traits. As in flashfm[4], the joint prior probability *p_i,j_^(1,2)^* for models *M_i_^(1)^*and *M_j_^(2)^* for traits 1 and 2, respectively, is defined as the product of the marginal prior probabilities when there is no model overlap of causal variants and is upweighted when there is sharing. That is, 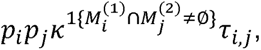 where ⎕ is a sharing parameter and *τ_i,j_* is a correction factor that guarantees that the prior probability of traits having particular model sizes is consistent for different values of ⎕; both parameters are derived in a combinatorial manner as in flashfm[4]. It follows that the trait-adjusted posterior probability for model *M_i_^(1)^*of trait 1 is calculated from

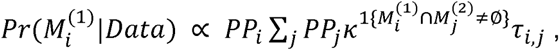

which is easily generalised to any number of traits *M* due to the traits being uncorrelated.

## Data and Code Availability

Summary statistics from the latent trait GWAS will be available from the GWAS Catalog. Custom code for the INTERVAL analyses are available at https://github.com/fz-cambridge/flashfmZero-INTERVAL-analysis.

FlashfmZero is freely available as an R package at https://jennasimit.github.io/flashfmZero[33]

## Acknowledgements

J.A. and F.Z. are supported by the UK Medical Research Council (MR/R021368/1 (J.A.), MC_UU_00002/4). W.J.A. is supported by NHS Blood and Transplant. A.B. was supported by core funding from the British Heart Foundation (RG/18/13/33946: RG/F/23/110103), NIHR Cambridge Biomedical Research Centre (NIHR203312) [*], BHF Chair Award (CH/12/2/29428) and by Health Data Research UK, which is funded by the UK Medical Research Council, Engineering and Physical Sciences Research Council, Economic and Social Research Council, Department of Health and Social Care (England), Chief Scientist Office of the Scottish Government Health and Social Care Directorates, Health and Social Care Research and Development Division (Welsh Government), Public Health Agency (Northern Ireland), British Heart Foundation and the Wellcome Trust.

Participants in the INTERVAL randomised controlled trial, who were recruited with the active collaboration of NHS Blood and Transplant England (www.nhsbt.nhs.uk), which has supported field work and other elements of the trial. DNA extraction and genotyping were co-funded by the National Institute for Health and Care Research (NIHR), the NIHR BioResource (http://bioresource.nihr.ac.uk) and the NIHR Cambridge Biomedical Research Centre (BRC-1215-20014) [*]. The academic coordinating centre for INTERVAL was supported by core funding from the: NIHR Blood and Transplant Research Unit (BTRU) in Donor Health and Genomics (NIHR BTRU-2014-10024), NIHR BTRU in Donor Health and Behaviour (NIHR203337), UK Medical Research Council (MR/L003120/1), British Heart Foundation (SP/09/002; RG/13/13/30194; RG/18/13/33946), NIHR Cambridge BRC (BRC-1215-20014; NIHR203312) [*], and by Health Data Research UK. A complete list of the investigators and contributors to the INTERVAL trial is provided in reference [12]. The academic coordinating centre would like to thank blood donor centre staff and blood donors for participating in the INTERVAL trial.

We thank Parsa Akbari for making available blood traits from the INTERVAL study adjusted for technical variation.

For the purpose of Open Access, the authors have applied a CC BY public copyright licence to any Author Accepted Manuscript version arising from this submission.

## Footnotes

* The views expressed are those of the authors and not necessarily those of the NIHR or the Department of Health and Social Care.

