## Supplemental Information for "Improved genetic discovery and fine-mapping resolution through multivariate latent factor analysis of high-dimensional traits"

**Figure S1.** **Basophil-related latent trait ML8 is associated with rs9310935, which has moderate signals with multiple basophil traits.** (A) regional association plot, highlighting rs9310935 and conditional regional association plot, conditioned on, rs3217673, rs163546, rs1669340, rs17027750, rs163563, rs3856850, rs334782, rs1695315, rs10212483, rs13097407, rs6787336 which are lead SNPs from previous publications. This is a supplement to Figure 3A.

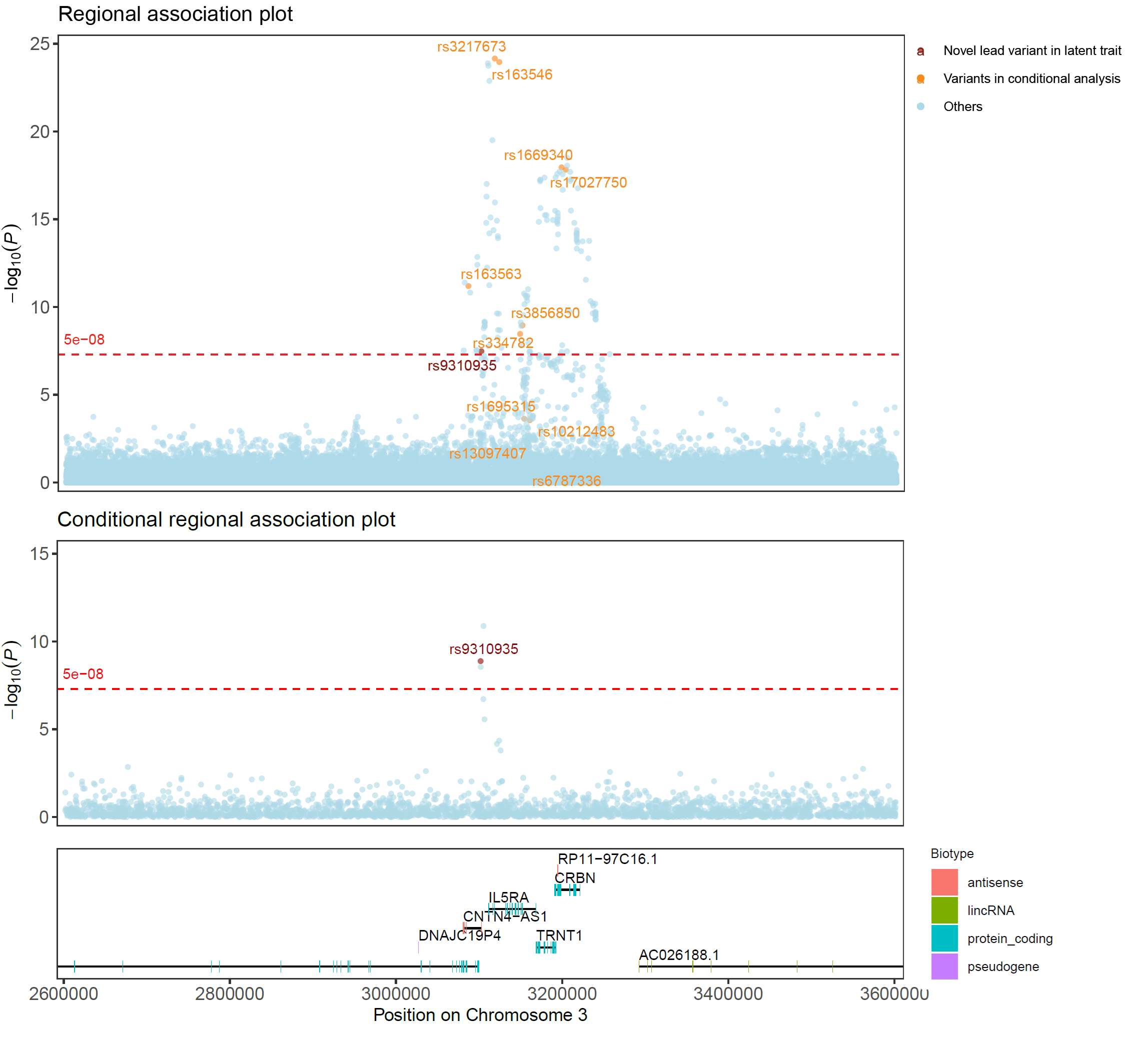

**Figure S2. Regional association plots of rs6064377 for ML6 and its related raw traits, HCT, RBC#, HGB, RPI.**  The top right panel also shows the ML6 association plot, conditioned on previously published lead variants.
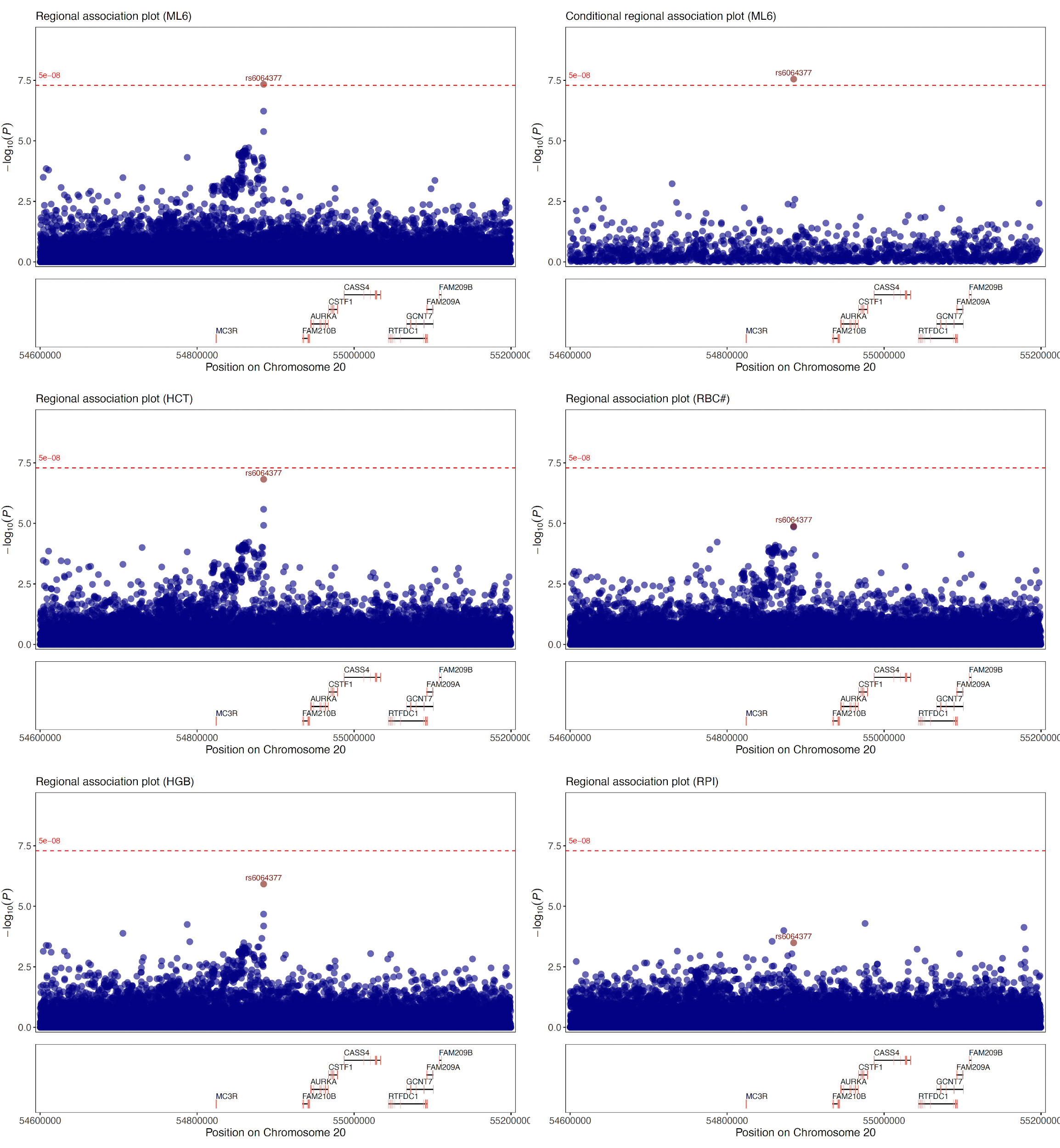

**Figure S3. Latent and raw trait correlations in the *PIEZO1* region show high correlations amongst each latent trait with their linked raw traits, and amongst raw traits having a common latent trait.** ML8 contributes to four basophil-related traits, ML12 contributes to MCHC, and ML13 contributes to three reticulocyte traits, as indicated by the correlation blocks.

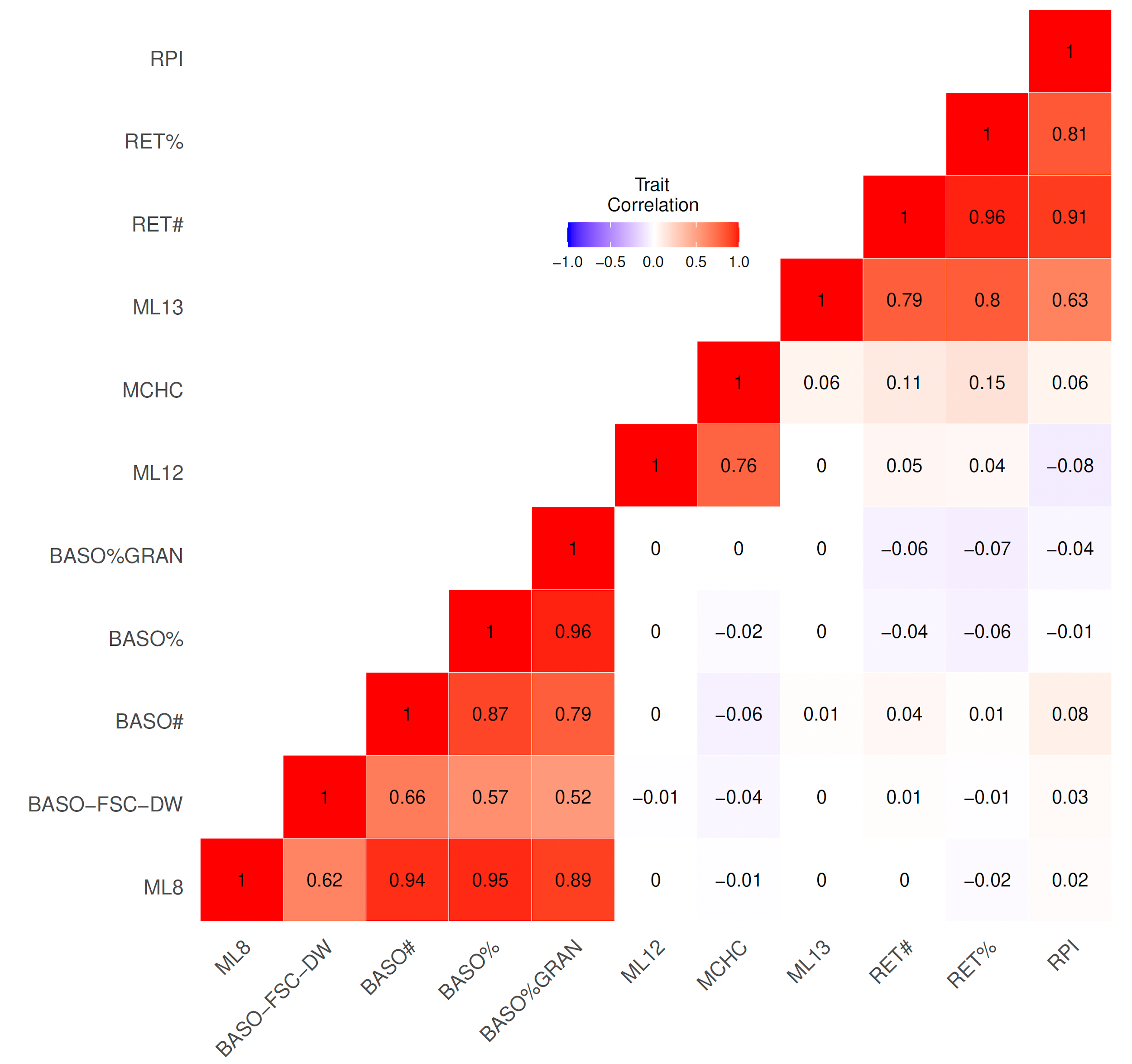

**Figure S4.** **In the *PIEZO1* region, the 99% credible sets (CS99) for raw traits become more refined for latent traits and gain further refinement by latent multi-trait fine-mapping.** Variants belonging to at least one CS99 are listed at the upper axis, labelled rsID. The CS99 for latent traits from single-trait fine-mapping are shown by the latent trait name, e.g. ML12. The CS99 for latent traits from multi-trait fine-mapping are shown by the latent trait name with an asterisk, e.g. ML12*. The coloured dots indicate which variants are members of the CS99 for the listed trait, where the size of each dot is proportional to the marginal posterior probability (MPP) of causality. Dot colours coincide with SNP groups, which are groups of variants in high LD (r^2^ > 0.8) and with MPP > 0.01 as estimated by the fine-mapping method. Boxes indicate latent and raw traits that are related to each other and the CS99 sizes are provided for each trait, listed by increasing size, e.g. ML12 and ML12* both have a CS99 of size 3 and MCHC has 4 variants in its CS99 - these three traits are related to each other as indicated by the box.

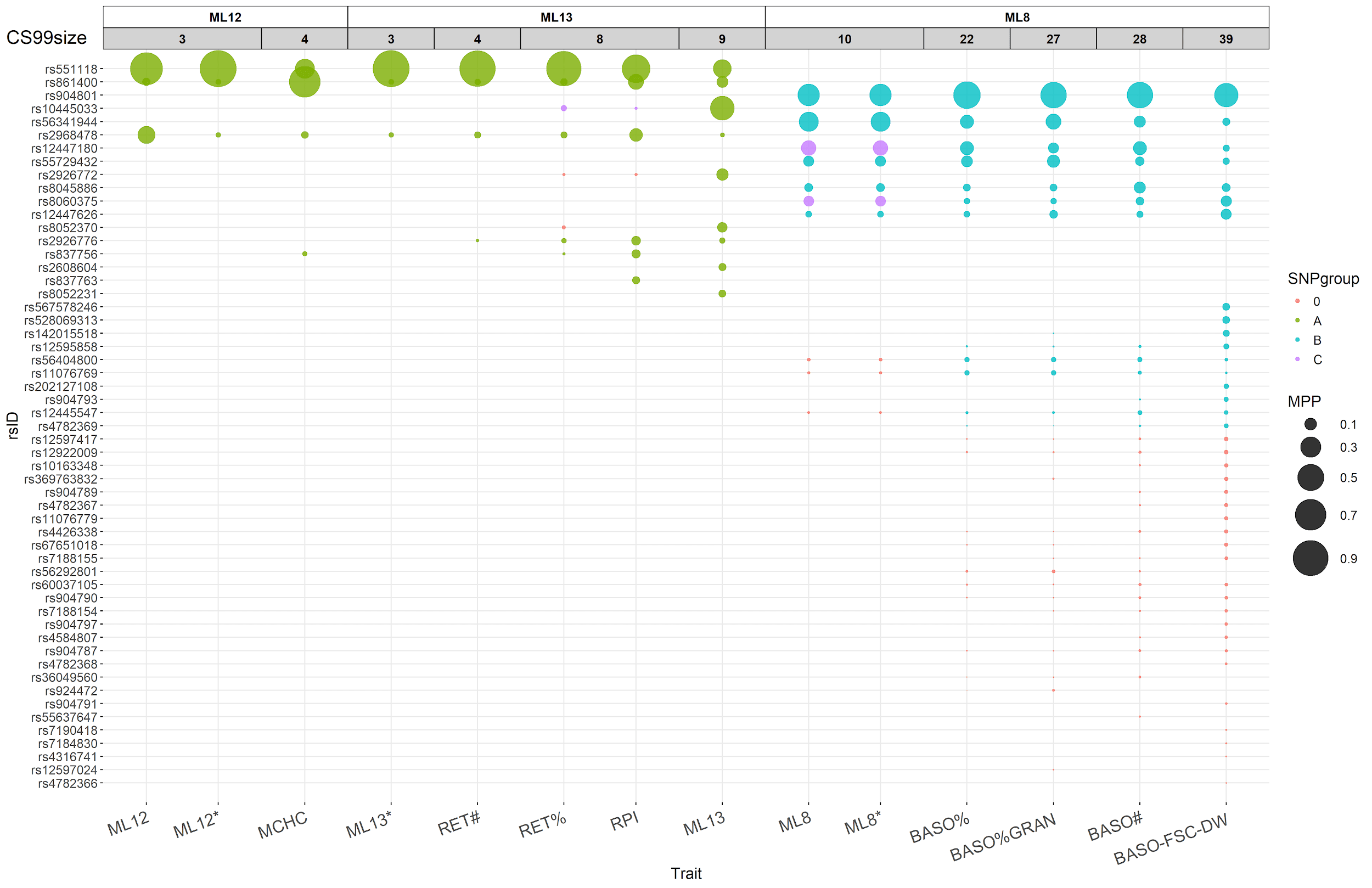

**Figure S5. Latent and raw trait correlations in the *TMCC2* region show high correlations amongst each latent trait with their linked raw traits, and amongst raw traits having a common latent trait.** ML5 contributes to nine platelet-related traits, and ML13 contributes to three basophil-related traits, as indicated by the correlation blocks.

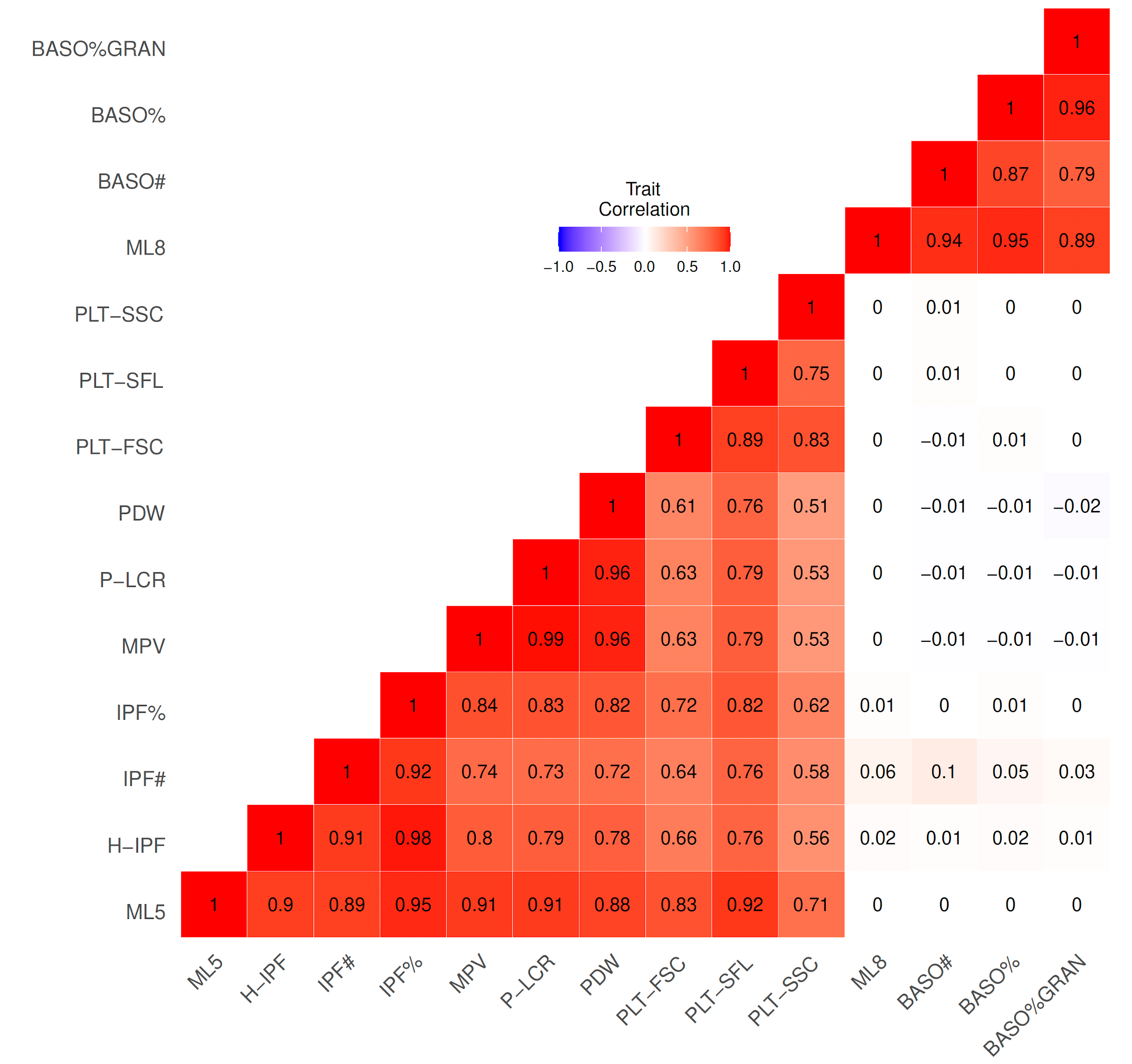

**Figure S6.** **In the *TMCC2* region, the 99% credible sets (CS99) for raw traits become more refined for latent single and multi-trait fine-mapping with identical results between single-trait and multi-trait fine-mapping because of no shared causal variants.** Variants belonging to at least one CS99 are listed at the left axis, labelled rsID. The CS99 for latent traits from single-trait fine-mapping are shown by the latent trait name, e.g. ML5. The CS99 for latent traits from multi-trait fine-mapping are shown by the latent trait name with an asterisk, e.g. ML5*. The coloured dots indicate which variants are members of the CS99 for the listed trait, where the size of each dot is proportional to the marginal posterior probability (MPP) of causality. Dot colours coincide with SNP groups, which are groups of variants in high LD (r^2^ > 0.8) and with MPP > 0.01 as estimated by the fine-mapping method. Boxes indicate latent and raw traits that are related to each other and the CS99 sizes are provided for each trait, listed by increasing size, e.g. ML8 and ML8* both have a CS99 of size 12, BASO# and BASO% both have CS99 sizes of 18, and BASO%GRAN has 19 variants in its CS99 - these five traits are related to each other as indicated by the box.

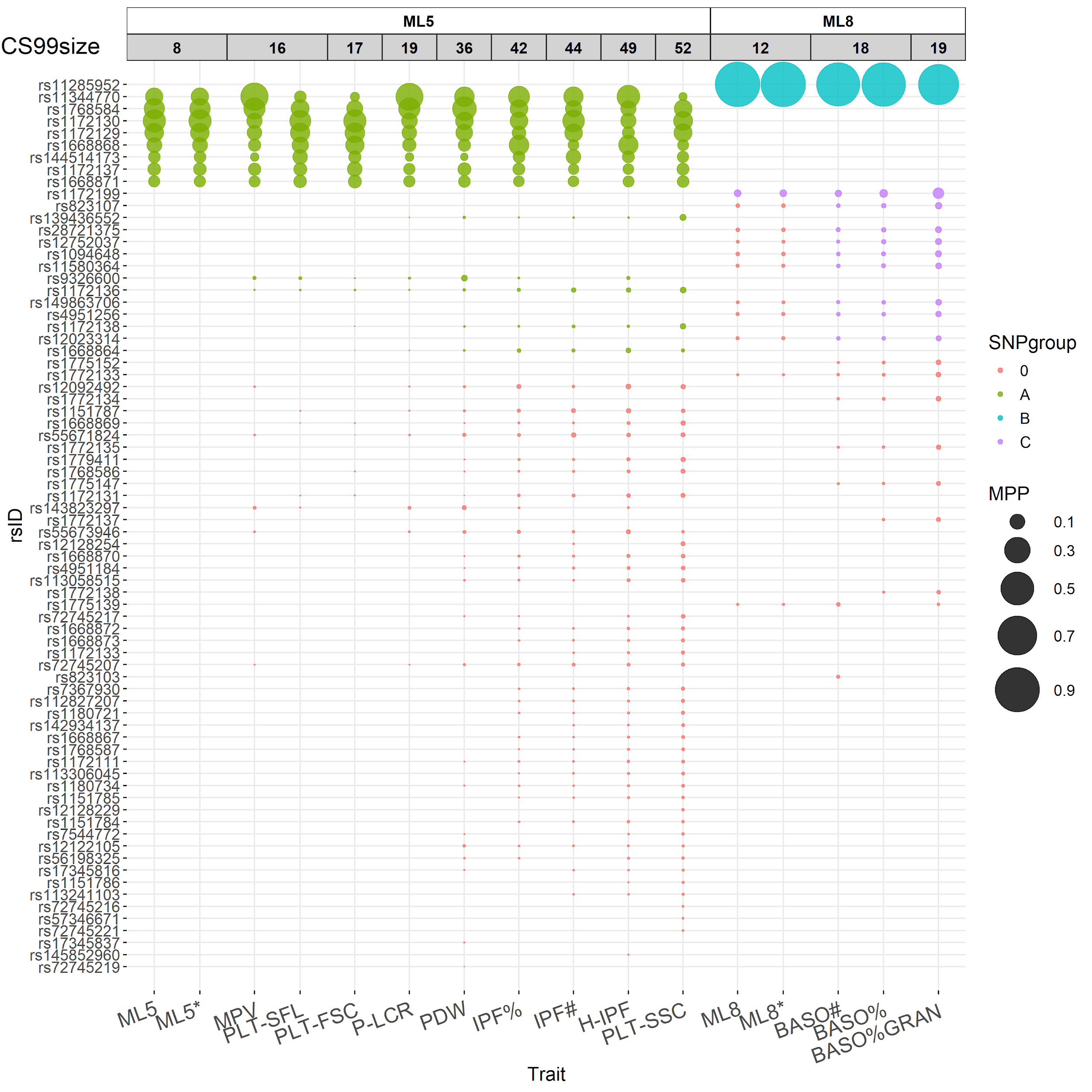
